# Morphogen-guided neocortical organoids recapitulate regional areal identity and model neurodevelopmental disorder pathology

**DOI:** 10.1101/2025.09.02.672952

**Authors:** Yuan-Chen Tsai, Hajime Ozaki, Xinyi Wang, Axel A. Almet, Isabel Fleming, Kaori Shiraiwa, Matthew Jung Min Noh, Caihao Nie, Sunnyana Trejo, Bret Kiyoshi Sugita, Jiya Dalal, Ruben Alberto Gonzalez, Briana De Jesus, Gregory Li-Min Chen, Michael J Gandal, Qing Nie, Momoko Watanabe

## Abstract

The human neocortex exhibits characteristic regional patterning (arealization) critical for higher-order cognitive function. Disrupted arealization is strongly implicated in neurodevelopmental disorders (NDDs), but current neocortical organoid models largely fail to recapitulate this patterning, limiting mechanistic understanding. Here, we establish a straightforward method for generating arealized organoids through short-term early exposure to anterior (FGF8) or posterior (BMP4/CHIR-99021) morphogens. These treatments created distinct anterior and posterior signaling centers, supporting long-lasting polarization, which we validated with single-cell RNA sequencing that revealed area-specific molecular signatures matching prenatal human cortex. To demonstrate the utility of this platform, we modeled Fragile X Syndrome (FXS) in organoids with distinct anterior and posterior regional identities. FXS organoids showed highly disrupted SOX4/SOX11 expression gradients along the anterior-posterior axis, consistent with alterations found in autism spectrum disorder (ASD) and demonstrate how regional patterning defects may contribute to NDD pathology. Together, our study provides a robust platform for generating neocortical organoids with anterior-posterior molecular signatures and highlights the importance of modeling NDDs using experimental platforms with neuroanatomic specificity.

## Introduction

Neocortical organoids derived from human pluripotent stem cells (hPSCs) are an invaluable tool for studying human neurodevelopment and disease. Organoids consistently self-organize to recapitulate the laminar architecture of the developing human cerebral cortex^1–4^. However, current methods^5–7^ of generating neocortical organoids still struggle to induce robust anteroposterior (AP) polarity that precedes ‘arealization’^8^ – the reproducible and stereotyped parcellation of the neocortex into areas serving distinct modalities, each with unique connectivity, cell types, cytoarchitecture, and gene expression signatures. This organization is critical for efficient neural processing and circuit formation and underlies much of human cognition and behavior^9^. In vivo arealization is initially guided by morphogens—extracellular signaling molecules secreted by localized organizers—that form gradients essential for cell specification, diversity, and topographical organization^8–10^. Two morphogen signaling pathways, Fibroblast Growth Factor (FGF) and Bone Morphogenetic Protein (BMP), antagonistically regulate areal identities along the AP axis by impacting downstream transcription factors^8, 10, 11^. FGFs are secreted by the anterior neural ridge (ANR), whereas BMPs and Wingless/Integrated (WNTs) are secreted from the roof plate, which later becomes the secondary organizer called the dorsomidline telencephalon (DMT), including the cortical hem, choroid plexus, and choroid plaque^10^.

In the developing human neocortex, areal signatures begin to emerge during the second trimester and display characteristic expression patterns in a cell-type-specific and time-dependent manner^12–15^. Most neocortical areas lack distinct transcriptional boundaries; instead, gene expression gradients—shaped by FGF and BMP signaling—underlie neocortical polarization^12–15^. These neocortical gene expression gradients are long-lasting, highly consistent and reproducible in the adult human brain across individuals^16^. Further, transcriptomic patterning of the neocortex has been associated with fMRI-based measures of resting state connectivity, indicating a potential functional link with neural circuitry^17^. Finally, transcriptomic areal identity is notably disrupted in NDDs such as ASD, which exhibits a broad attenuation of typical transcriptomic gradients across the neocortex with posteriorly patterned regions exhibiting the greatest disruption^18^.

As neocortical organoids lack external (e.g., sensory) inputs, they offer a unique model to test the “protomap hypothesis” – namely, whether the neocortex is intrinsically pre-patterned to generate distinct areal signatures or whether this process requires inputs from other brain regions. Both FGF8 and retinoic acid promote anterior regional signatures in neocortical organoids, but with limited axial polarity defined by a few transcriptional markers^2, 15, 19^. An assembloid-based approach to anteriorize organoids has also been reported^20^. However, methods to develop neocortical organoids with specific posterior areal identities have not yet -- to our knowledge -- been reported^2, 15, 19, 20^. Here, we establish a simple method to induce polarization in neocortical organoids via short-term FGF8 or BMP4/CHIR-99021 (canonical WNT agonist) exposure, generating homeodomain protein gradients and AP identities in a cell-type– and time-dependent manner. Immunohistochemistry and single-cell RNA sequencing (scRNA-seq) revealed self-polarizing organizers and graded areal signatures. Applying this approach to patient-derived hiPSCs from individuals with Fragile X Syndrome (FXS) revealed disrupted SOX4 and SOX11 gradients, mirroring transcriptomic findings observed in ASD patient tissue^18^. This platform offers a tractable system for investigating intrinsic gene regulatory networks and studying area-specific molecular pathology associated with NDDs.

## Results

### Neocortical organoids with temporally dynamic laminar specificity

To generate neocortical organoids, we followed our previously established approach adapted from the improved, serum-free floating culture of embryoid body-like aggregates with quick reaggregation (SFEBq; Methods), as summarized in Figure S1A^2, 4, 21^. By week (W) 2.5, the organoids express markers of neocortical progenitors (COUP-TF1, EMX1/2, FOXG1, PAX6, SOX2), apical (NCAD, aPKC), and basement (LAMININ) membranes (Figures 1A-B and S1C-S1D’), resembling neural tube formation at human gestational week (GW) 3-4^22^. By W5, the neuroepithelial cell (NEC) layer has rolled over and formed neural rosettes (Figure 1B)^2, 4, 21^. Distinct neocortical layers are now apparent, including the ventricular zone (VZ) with SOX2⁺PAX6⁺COUPTF1⁺SP8⁺ apical radial glial cells (aRGCs); the subventricular zone (SVZ) with TBR2⁺ intermediate progenitors (IPs); and the preplate (PP)/Cajal-Retzius (CR) region with TBR1⁺CTIP2⁺RELN⁺ neurons (Figures 1B, S1E–S1E’). The formation of these laminar structures mimics that of the developing human neocortex at GW5-7^23^. By W8, the formation of the cortical plate (CP) with TBR1⁺CTIP2⁺ neurons and the subplate (SP), strongly CSPG-positive^2, 23^, have occurred (Figure 1B).

**Figure 1:**
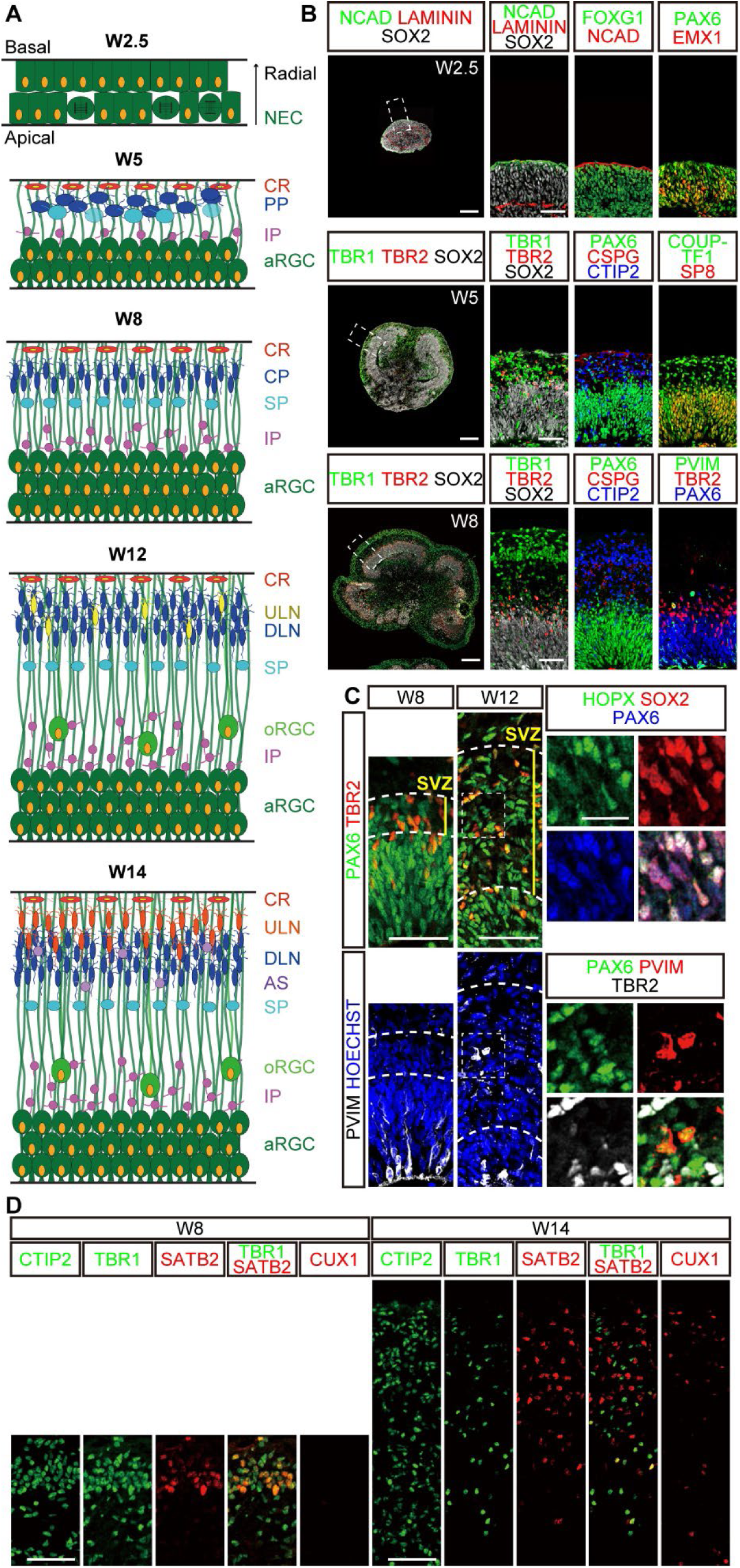
Neocortical organoids with temporally dynamic laminar layering. (A) Timeline of neocortical development from W2.5 to W14, showing key cell types. NEC: neuroepithelial cells; CR: Cajal Retzius neurons; PP: preplate neurons; IP: intermediate progenitors; aRGC: apical radial glial cells; CP: cortical plate neurons; SP: subplate neurons; ULN: upper-layer neurons; DLN: deep-layer neurons; oRGC: outer radial glial cells; AS: astrocytes. (B) Immunostaining of organoids at W2.5, W5, and W8 using cell-type markers. Left: whole organoids. Right: cropped regions (white boxes). Scale bars: 200 µm (left), 50 µm (right). (C) The subventricular zone (SVZ) expands from W8 to W12. Left: SVZ region with IPs and oRGCs. Right: zoom-ins (white boxes). Scale bars: 50 µm (left), 20 µm (right). (D) ULN and DLN markers in W8 and W14. DLN: CTIP2+, TBR1+. ULN: SATB2+, CUX1+. Scale bars: 50 µm.

After W8, organoids continue to develop and exhibit human-like features. By W12, the SVZ has expanded with abundant outer radial glial cells (oRGCs; PAX6⁺SOX2⁺HOPX⁺pVIM⁺TBR2⁻), one of the salient features of the neocortex in humans compared to rodents (Figure 1C)^24^. By W14, we observe CTIP2^+^TBR1^+^ deep-layer neuron-like cells (DLN) and SATB2^+^CUX1^+^ upper-layer neuron-like cells (ULN)^25^ (Figure 1D). Furthermore, SATB2^+^CUX1^+^ ULNs are clearly located superficially in the CP, recapitulating the inside-out layering of the GW15 human prenatal neocortex^4^. We also confirmed the presence of GFAP^+^HepaCAM^+^ astrocytes (Figure S1F-S1F’), which are also formed by mid-gestation *in vivo*^4^. Together, these data demonstrate that our neocortical organoids recapitulate the temporal progression of laminar structuring and cell type specification, spanning from embryonic timepoints to the mid-second trimester^4^, when transcriptomic areal signatures emerge^12–15^.

### Polarization of the neocortex with the induction of organizers

To generate anterior- or posteriorly polarized neocortical tissue, we added rostral or caudal morphogens from 15 to 21 days of differentiation, corresponding to the formation of primitive NECs and in vivo organizers^26^ (Figures 2A and 2C). We evaluated tissue polarization at W5 via expression of the canonical anterior and posterior homeodomain transcription factors SP8 and COUP-TF1 (Figures 1A and 2A-2C). Without growth factors, ∼50% of cells co-expressed both COUP-TF1 and SP8, resembling the parietal/temporal/occipital lobes of the GW10-14 human neocortex (Figures 2B and 2D). The extensive overlap of SP8/COUP-TF1 expression in the intermediate regions of the neocortex is unique to humans and not found in the mouse neocortex^27, 28^. Human neocortical organoids can also replicate this distinctive feature.

**Figure 2:**
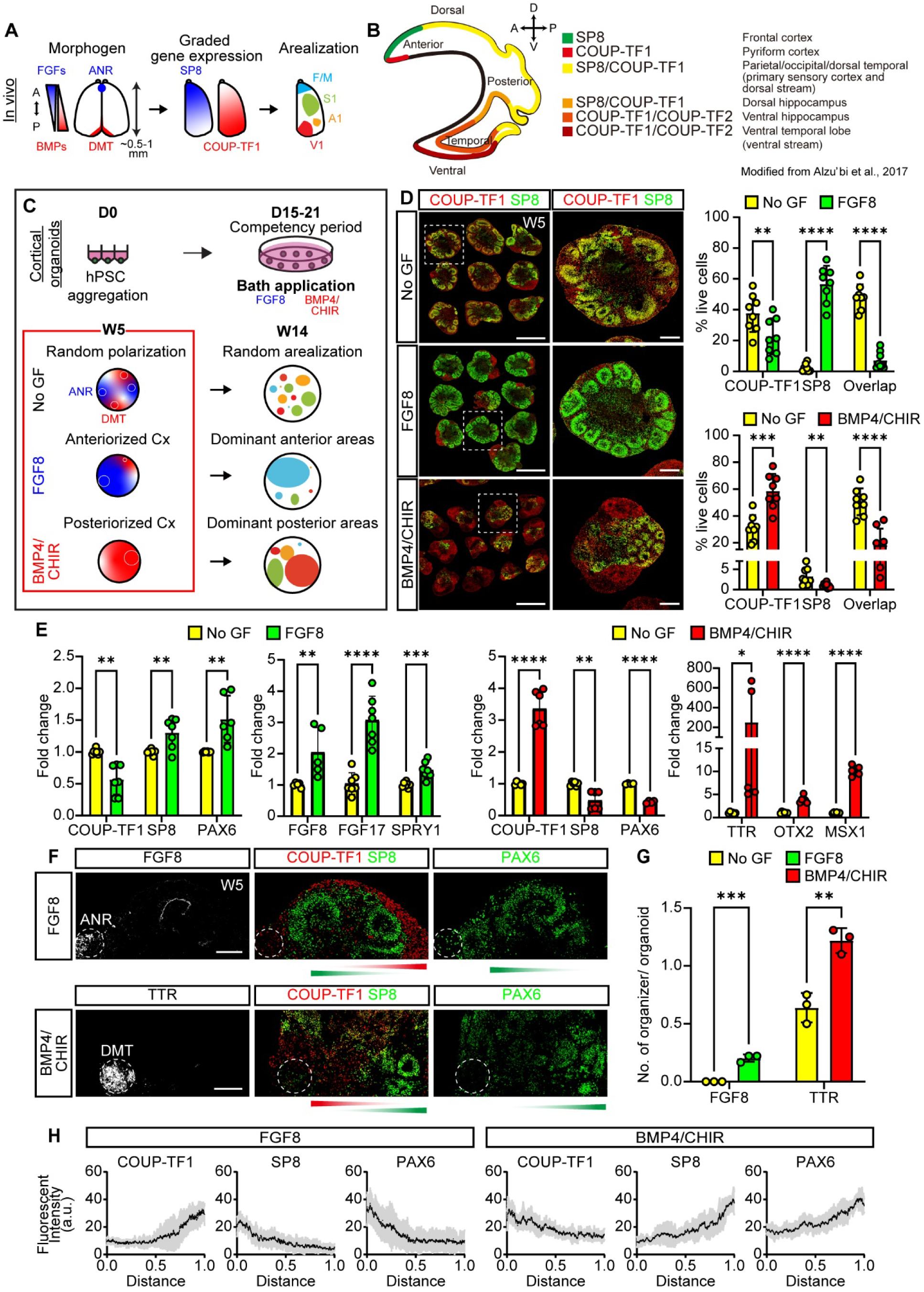
Polarization of the neocortical tissue with the induction of organizers. (A) Illustration of morphogen secretion from anterior (ANR) and posterior (DMT) organizers, leading to the polarization, graded gene expression, and arealization of the neocortex. Primary area. F/M: prefrontal/motor; S1: somatosensory; A1: auditory; V1: visual areas. (B) Sagittal view of SP8 and COUP-TF1 expression in W8 human fetal telencephalon, adapted from Alzu’bi et al 2017^16^. (C) Experimental paradigm for neocortical organoids with no growth factor (GF), FGF8, or BMP4/CHIR-99021. (D) Immunostaining for SP8 (anterior) and COUP-TF1 (posterior) in treated organoids. Left panels: tiled images. Middle: representative organoids. Right: quantification of COUP-TF1⁺, SP8⁺, and double-positive cells (N = 8 organoids, 3 batches per condition). Scale bars: 1000 µm (left), 200 µm (middle). (E) RT-qPCR for anterior/posterior transcription factors and signaling response genes (N = 4–7 samples; each pooled from 4–6 organoids; 3 batches). Markers: SP8, PAX6 (anterior); FGF8, FGF17 (anterior morphogens); SPRY1 (FGF-response gene); COUP-TF1 (posterior); TTR, OTX2 (DMT); MSX1 (BMP-response gene). (F) Induction of FGF8^+^ ANR and TTR^+^ DMT in FGF8- and BMP4/CHIR-treated organoids, respectively. Scale bar: 100 µm. (G) Quantification of organoids with ANR or DMT across conditions (N = 3 batches; 3 organoids/condition). (H) Normalized fluorescent intensity profiles of SP8, COUP-TF1, and PAX6 across organoid diameter (N = 7–8 organoids, 3 batches/condition). Black line: mean; grey lines: individual organoids. Data are presented as mean ± SD in D, E, and G. P-values calculated by unpaired t-Test. *p<0.05; **p<0.01; ***p<0.001; ****p<0.0001.

To anteriorize neocortical organoids, we added 400 ng/mL FGF8 (see titration assay; Figures S2A and S2B), leading to strong upregulation of the anterior marker SP8 (∼55% SP8^+^ single-positive cells). In contrast, double-positive COUP-TF1/SP8 cells and COUP-TF1 single-positive cells decreased to 5 % and 20 %, respectively (Figure 2D). The increase in single-positive SP8 cells in FGF8-treated organoids was comparable to that of GW10-14 human frontal lobe^27^ (Figure 2B). FGF8 application upregulated anterior (SP8 and PAX6) and FGF8 target genes (FGF8, FGF17, and SPRY1) (Figure 2E). Given its key role in primate prefrontal cortex (PFC) development^15, 19^, we tested whether treatment with retinoic acids would further enhance the induction of the anterior identity. From W5, organoids were cultured with or without the retinoic acid precursor Vitamin A (Figures S2C-S2F). Consistent with anterior identity, we observed upregulation of the PFC marker CBLN2 and the motor cortex marker CYP26B1 (Figure S2F). Either vitamin A alone, or in combination with FGF8, led to a decrease in COUP-TF1 expression and an increase in COUP-TF1^+^SP8^+^ cells (Figures S2D and S2E), but we did not observe any additive effect. These results indicate that both FGF8 and retinoic acids promote anterior neocortical identity.

To generate neocortical organoids with a posterior areal identity, 50 ng/mL BMP4 and 3 μM CHIR-99021 (WNT activator) were added (see titration assays; Figures 2D and S3A). The expression of COUP-TF1 was significantly increased, whereas SP8 and PAX6 were downregulated (Figures 2D and S3A-S3B). SP8-single positive cells were rare, while about 20% of cells were double positive for COUP-TF1 and SP8 (Figure 2D), recapitulating parietal/temporal/occipital lobes *in vivo*^27^ (Figure 2B). The increase of COUP-TF1 single-positive cells likely reflects the induction of posterior regions, including the hippocampal primordium (Figures 2B, 2D, and S3A-S3E), as previously characterized^4, 29^.

To further establish that FGF8 and BMP4/CHIR treatment induced AP axis arealization in neocortical organoids, we investigated the induction of organizers in these conditions. FGF8 treatment resulted in increased FGF8⁺ ANR-like cells (Figures 2F, 2G, and S3C; average of 0.2 organizers/organoid section), which were located adjacent to regions with high expression of SP8 and PAX6 and low expression of COUP-TF1 (Figure 2H). On the other hand, BMP4/CHIR application led to increased TTR^+^ choroid plexus epithelial cells, posterior organizers (Figures 2F and 2G; average of 1.2 organizer/organoid section; see Figure S3B, S3D-S3E for additional markers OTX2, and MSX1), and the adjacent areas exhibited high expression of COUP-TF1 and low expression SP8 and PAX6 (Figure 2H). Hence, the short-term application of FGF8 and BMP4/CHIR induced AP organizers in neocortical organoids, which in turn likely self-polarize into anterior or posterior areas.

### Area-specific transcriptional signatures in polarized neocortical organoids

Neocortical areas are initially defined by cell-type-specific and temporally dynamic gene expression gradients, rather than exclusive gene expression sets^5, 12–15, 30^. Thus, we performed scRNA-seq (Figure 3A) to investigate whether polarized organoids recapitulate the molecular signatures associated with distinct neocortical lobes/areas^12, 14, 15, 20^. We characterized key molecular features at W5 (PP stage), W8 (CP stage), and W14 (ULN emergence) using neocortical organoids with and without the application of FGF8 or BMP4/CHIR-99021.

**Figure 3:**
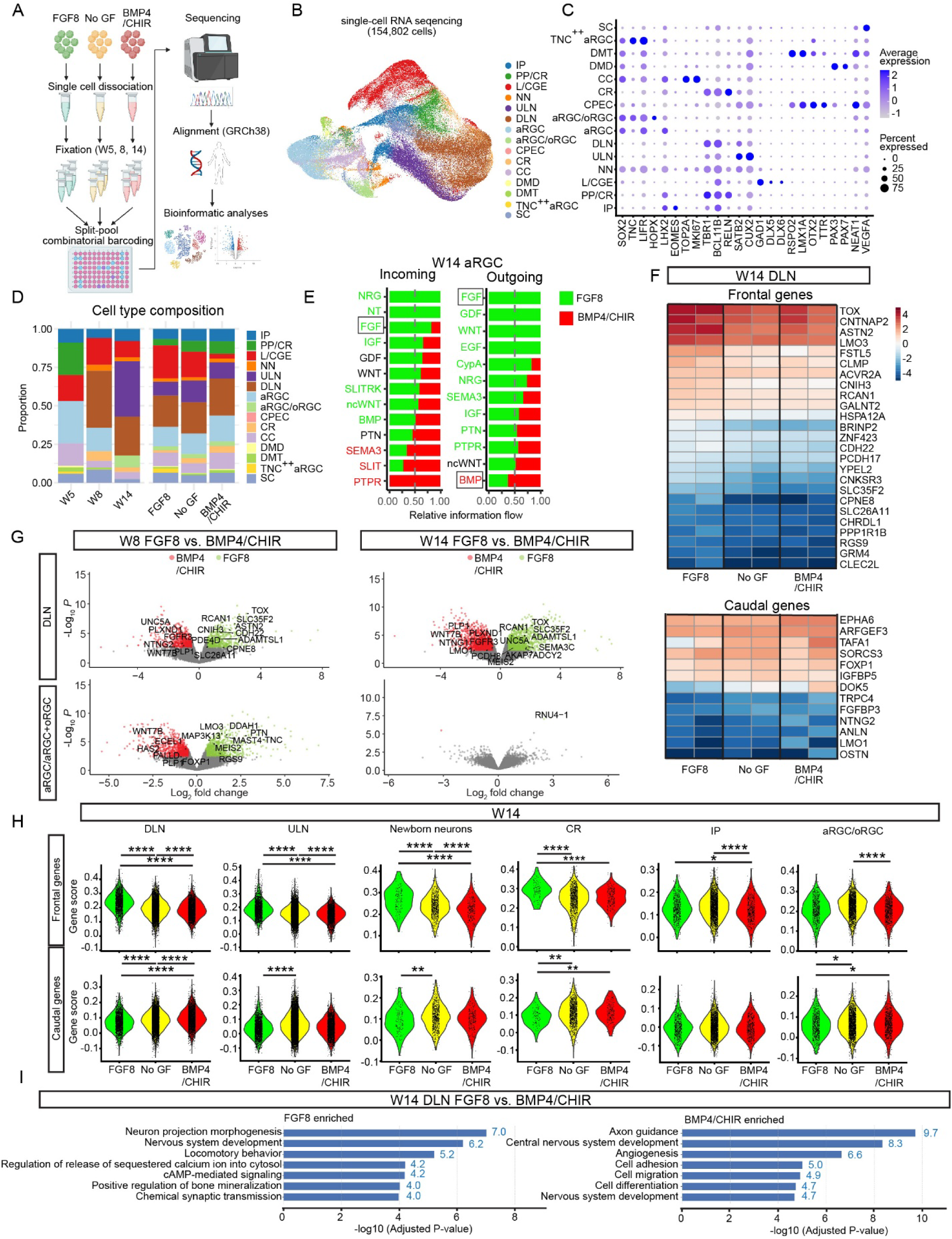
Areal transcriptional signatures in polarized neocortical organoids. (A) Sample processing workflow. (B) UMAP of 154,802 cells (post-QC) across all timepoints and conditions, revealing 15 clusters. IP: intermediate progenitors; PP/CR: preplate/Cajal Retzius neurons; L/CGE: lateral/caudal ganglionic eminence; NN: newborn neurons; ULN: upper-layer neurons; DLN: deep-layer neurons; aRGC: apical radial glial cells; oRGC: outer radial glial cells; CPEC: choroid plexus epithelial cells; CC: cycling cells; DMD: dorsomidline diencephalon; DMT: dorsomidline telencephalon; SC: stressed cells. (C) Dot plot showing mean expression (color) and proportion (size) of canonical markers across clusters. (D) Cell type composition by timepoint (left) and condition (right). (E) CellChat analysis for W14 aRGC/oRGC for incoming and outgoing signals. (F) Expression of frontal (top) and caudal (bottom) genes in W14 DLN. (G) Volcano plots of differentially expressed genes in DLN and aRGC/oRGC clusters at W8 and W14 under FGF8 vs BMP4/CHIR. Colored: genes with false discovery rate FDR<0.05 and ⏐log2FC⏐>0.5. (H) Gene score analysis for known frontal (upper panel) and caudal area genes (lower panel) for neocortical cell types at W14. See Table S5 for the full statistical comparisons. Statistics: Tukey’s Honest Significance Distance test. *p<0.05; **p<0.01; ***p<0.001; ****p<0.0001. (I) Gene ontology analysis for biological processes of DEGs in FGF8 vs BMP4/CHIR-treated W14 DLN.

Following dimensionality reduction and clustering, we identified 15 cellular clusters expressing canonical cell-type markers, including aRGC, oRGC, IP, CR, DLN, and ULN (Figures 3B and 3C; Table S1). All cells broadly expressed the forebrain marker FOXG1 (Figure S4D). All major regions of the forebrain, including the cerebral cortex (LHX2^+^EMX2^+^TBR1^+^PAX6^+^), dorsomidline telencephalon (DMT: RSPO2^+^LMX1A^+^OTX2^+^TTR^+^), dorsomidline diencephalon (DMD: PAX3^+^PAX7^+^), and lateral and caudal ganglionic eminences (L/CGE-like cells: GAD1^+^DLX5^+^DLX6^+^) were identified (Figures 3B-C, and S4D). We observed that developmental timepoint was the primary source of segregation (Figure S4A), as reported in the developing mouse brain^31^. Finally, cell types were highly overlapping across all three experimental conditions (no morphogen, FGF8, or BMP4/CHIR; Figure S4B).

Cell type composition analyses (Figure 3D) revealed temporally dynamic patterns of cell-type formation, following typical developmental progressions, as seen in immunochemical analyses (Figure 1). Cell type composition was similar across conditions (Figures 3D and S4B), although some expected differences were noted across the dorsal-ventral axis. TNC^++^ aRGCs cluster was only detected in W5 and W8 FGF8-treated organoids (Figures 3D and S4F), which showed high expression of the ventral marker DLX2 and lower expression of the neocortical marker EMX2, consistent with its role in induction of the ganglion eminence^32–34^. Application of BMP4/CHIR, which dorsalizes the entire telencephalon, therefore was associated with reduced formation of ventral L/CGE-like cells (Figure 3E).

To further understand the effects of FGF8 and BMP4/CHIR on neocortical AP axis arealization, we next conducted differential expression analyses (Methods) comparing morphogen effects on cell types with neocortical identities (EMX2^+^LHX2^+^TBR1^+^), including: DLN, ULN, NN, CR, IP, aRGC, and aRGC/oRGC. Differentially expressed genes (DEGs) were temporally dynamic and highly distinct across cell-types (Figure S4G). Primary progenitors (aRGC/oRGC) exhibited more DEGs at early (W5, W8) compared to later stages (W14), whereas secondary progenitors (IP) had consistently high and increasing numbers of DEGs over time. We observed fewer DEGs within ULN, perhaps due to its relatively late emergence at W14. To assess whether the polarization persisted at later stages, we conducted CellChat^35^ analysis on W14 aRGC/oRGC (Figure 3E). FGF and BMP pathways were found in FGF8 and BMP4/CHIR conditions, respectively, for both incoming and outgoing signals. This result suggests that brief early exposure to FGF8 and BMP4/CHIR treatments has lasting effects on molecular pathways in primary progenitors, likely due to the organizer formation (Figures 2F and 2G) that sustains neocortical organoid self-polarization.

We next compiled gene signatures (Table S6, Figure S6A) of spatially segregated neocortical regions, including anterior/frontal (frontal lobes/PFC) and posterior (temporal/occipital/visual areas) areas, leveraging data from scRNA-seq studies profiling human brain development from Shibata et al, 2021^15^, Bhaduri et al, 2021^12^, Qian et al, 2025^14^, and Bosone et al, 2024^20^. We examined the overlap between gene expression signatures of these anterior and posterior brain regions in the DEGs of FGF8, no GF, and BMP4/CHIR. At W14, frontal region-specific genes were enriched in the FGF8 group, whereas posterior-enriched genes were elevated in the BMP4/CHIR group (Figures 3F, 3G, and S5A). Both DLN and IP had large numbers of FGF8- or BMP4/CHIR-enriched genes associated with frontal and posterior regional signatures in W8 and W14 (Table S4). In contrast, the aRGC/oRGC population displayed limited frontal and posterior DEGs at W14, probably due to fewer primary progenitor cells.

To validate scRNA-seq DEG results, we next conducted immunochemical analyses of key morphogen-induced genes that exhibited consistent effects across time points and cell types (Table S4) and elevated in expected anterior or posterior regions of the developing mouse neocortex (Figures S6). For frontal areal signatures, these genes included CBLN2, TOX, and TNC (Figures S6B-S6D). Consistent with previous literature^14, 15^, CBLN2 expression was upregulated specifically in ULN (SATB2^+^), but not in DLN (CTIP2^+^) (Figure S6G). In agreement with bioinformatic analyses, TOX expression was elevated in CTIP2⁺ neurons in FGF8-treated organoids (Figure S6C). Higher TNC fluorescence was detected within SOX2^+^ aRGC progenitor regions in the FGF8 condition (Figure S6D). For posterior areal signatures, we found FOXP1 and LMO1 were consistently upregulated in aRGCs or neurons, respectively, following BMP4/CHIR treatment (Figures S6E and S6F). Finally, we performed Foxp1 immunostaining in E12.5 mouse brain tissue and confirmed the enriched Foxp1 expression in the posterior neocortex (Figure S6H). Together, bioinformatically predicted frontal and posterior areal-specific genes were validated in polarized neocortical organoids, indicating transcriptional recapitulation of initial neocortical arealization.

Finally, we developed a “areal gene score” (see Methods) representing the aggregated expression of frontal and posterior gene sets^12, 14, 15, 20^. Frontal gene scores showed a graded decrease from FGF8 to no-GF to BMP4/CHIR-treated organoids in a cell-specific and time-dependent manner. Posterior gene scores showed the opposite trend. In accordance with human fetal cortex findings, graded gene scores were prominent in neurons but less pronounced in primary progenitors at W14 (Figures 3H and S4D-E). Gene ontology analyses indicated similar pathway enrichments for DEGs across FGF8 and BMP4/CHIR conditions, including nervous system development, neuron projection, axon guidance, cell adhesion, and migration (Figures 3I and S5B-C). Together, these results show that morphogen-specific arealization of neocortical organoids results in concordant patterns of frontal/anterior or posterior brain-region specific gene expression, thus recapitulating morphogen-induced intrinsic mechanisms of the neocortical arealization.

### Neocortical organoids with areal signatures for disease modeling

As a proof of principle, we investigated whether polarized neocortical organoids could serve as a modelling platform to study differential phenotypic effects on neocortical areas within the context of NDD. We first took the DEGs from the comparison of FGF8 and BMP4/CHIR conditions to examine overlap with known disease genes, including Fragile X Syndrome (FXS)^36^, high-confidence risk genes for ASD^37, 38^ or broad NDDs^38^. By W14, the DLN and IP showed 143 and 231 FMRP-interacting genes^36^, 201 and 312 ASD-associated genes^37^, and 1,800 and 2,668 NDD-associated genes^38^, respectively (Figure S7A), emphasizing the importance of using neocortical organoids with areal signatures in disease modeling.

Next, we established neocortical organoids generated from hiPSCs produced from the fibroblasts of patients with FXS, one of the most common inherited causes of intellectual disability and a syndromic cause of ASD, to model area-specific phenotypes. Previous work has indicated that both rare and common ASD-associated genetic variants converge, particularly within known FMRP target genes co-expressed during early human brain development^39^. Further, ASD is characterized by a highly consistent molecular pathology, with widespread transcriptional dysregulation observed across the neocortex^18^. We confirmed that FXS neocortical organoids did not have detectable *FMR1* and FMRP expression (Figures S7B-S7C)^40^. Before W14, FXS organoids did not exhibit a neural induction phenotype (Figures 4A and 4B). At W14, they exhibited more aRGCs and fewer DLNs, suggesting premature neuronal differentiation (Figure 4B). The attenuation of AP expression gradients of SOX4 and SOX11 was identified in patients with ASD (Figure 4C)^18^. Utilizing FXS organoids with and without FGF8 or BMP4/CHIR, we similarly found that SOX4 and SOX11 expression gradients were detected in neurotypical neocortical organoids, but not in FXS neocortical organoids (Figures 4D-4F). We confirmed that anteroposteriorization was present through other canonical transcription gradients (COUPTF1, SP8, and PAX6) in FXS neocortical organoids (Figure S7D). Together, neocortical organoids with frontal and caudal signatures are crucial for identifying differential phenotypic effects on neocortical areas in disease modeling.

**Figure 4:**
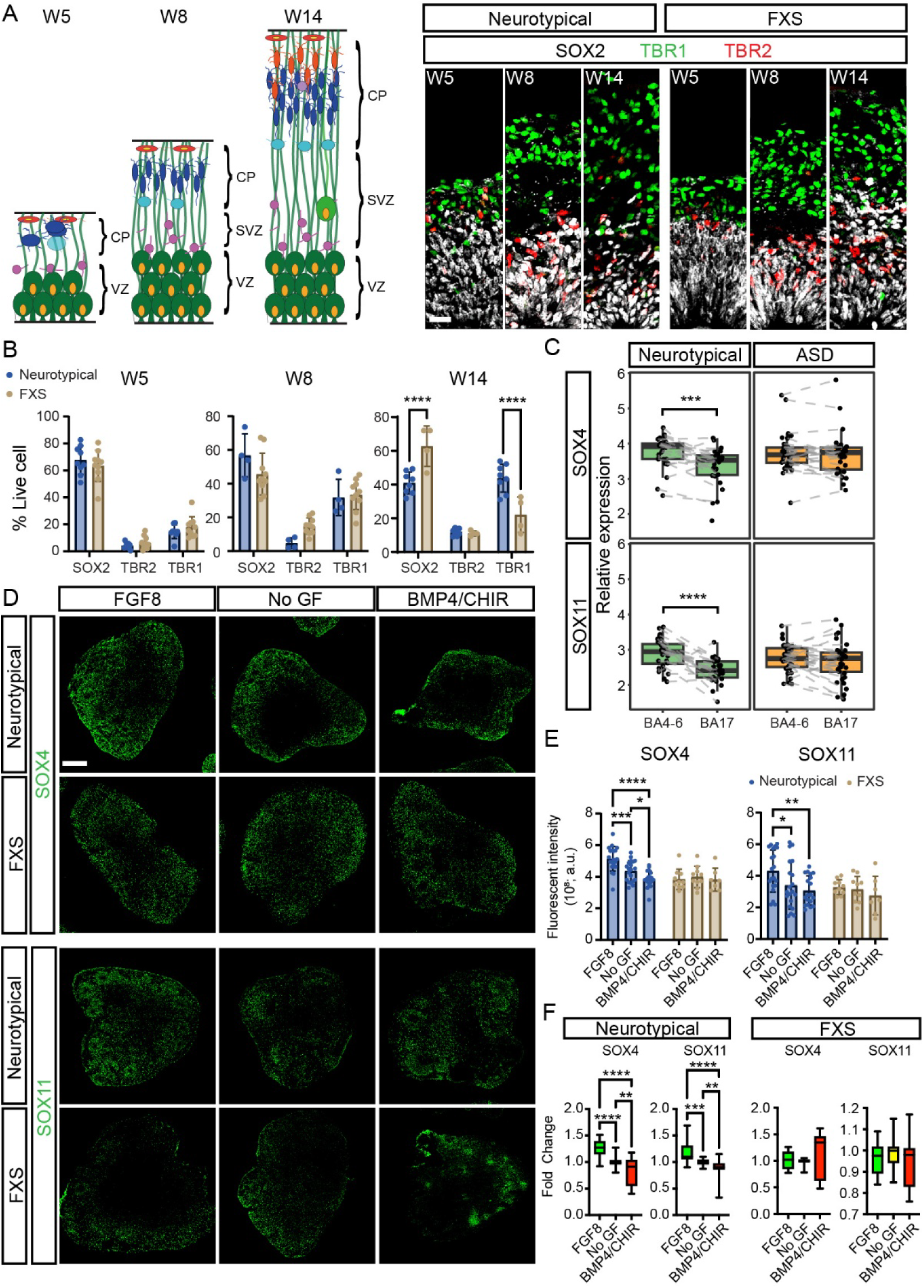
Fragile X Syndrome (FXS) disease modeling using neocortical organoids with areal signatures. (A) Cortical development in neurotypical and FXS organoids (W5–W14). Left: schematic of cortical layers. Right: immunostaining for SOX2 (neural progenitors), TBR2 (intermediate progenitors), and TBR1 (DLN). Scale bar: 20 µm. (B) Quantification of SOX2+, TBR2^+^ and TBR1^+^ cells in total live cells at W5, W8, and W14. N = 4-10 organoids/timepoint/cell line from 2–3 batches (3 cell lines total). (C) Human postmortem data from Gandal et al., 2022: SOX4 and SOX11 expression in frontal (BA4–6) vs. occipital (BA17) cortex in neurotypical vs. ASD patients. Statistics: paired Wilcoxon signed-rank test. (D) Immunostaining of SOX4 and SOX11 in W8 neurotypical or FXS organoids in FGF8, no GF, and BMP4/CHIR conditions. (E) Fluorescent quantification of SOX4 and SOX11 in whole organoids. “IntDen” values, the product of area and mean gray value, were used for comparisons. N = 7–10 organoids/condition from 3 cell lines. (F) RT-qPCR of SOX4 and SOX11 in neurotypical (N = 30–33, 2 cell lines, 2 batches/condition) and FXS (N = 11–13, 1 cell line, 4 batches/condition) organoids across conditions. Statistics: Two-way ANOVA with post-hoc test Tukey in B and E. One-way ANOVA with post hoc test Tukey in F. All data are represented as mean ± SD. *p<0.05; **p<0.01; ***p<0.001; ****p<0.0001.

## Discussion

This study establishes a straightforward, accessible approach to generate polarized neocortical organoids with stable frontal and posterior areal signatures, using short-term FGF8 or BMP4/CHIR exposure. Anterior and posterior organizers emerged by W5, likely contributing to continued polarization. We utilized scRNA-seq and bioinformatic analysis to quantify graded gene expression of areal signatures, which has been challenging to study because it is temporally dynamic and cell-type specific^12–15, 30^. In addition to well-known frontal markers (e.g., CBLN2), we identified TNC, TOX, FOXP1, and LMO1 as consistent and reliable areal signatures. Lastly, FXS organoids showed area-specific molecular alterations, highlighting the utility of polarized organoids for modeling area-specific disease features. In summary, the simple application of FGF8 or BMP4/CHIR provides a practical framework for producing polarized neocortical organoids suited for modeling spatially distinct neurodevelopmental disorders.

A long-standing question in neurodevelopment is whether neocortical arealization is driven primarily by intrinsic or extrinsic mechanisms^8, 10, 26^. The ‘protomap’ model focuses on intrinsic factors and posits that neocortical areas develop without external inputs. On the other hand, the ‘protocortex’ model highlights the importance of extrinsic influences, proposing that afferent projections shape neocortical area identity. Neocortical organoids with distinct areal identities offer a unique opportunity to dissect intrinsic and extrinsic mechanisms of arealization in a controlled system.

In this paper, Cajal Retzius (CR) cells also exhibited graded expression patterns of frontal and caudal genes in neocortical organoids with or without FGF8 or BMP4/CHIR (Figure 3H). CR cells are among the earliest-born neurons in the forebrain and arise from distinct sources, including the cortical hem, anti-hem, septum, and thalamic eminence^41, 42^. Because BMP/CHIR application increases the presence of the caudal cortical hem (Figure S3) in organoids, they might reflect different developmental origins, and how that would contribute to the areal differences in the laminar organization might be an interesting topic to pursue.

### Limitations of the study

In this study, we optimized FGF8 and BMP4/CHIR-99021 concentrations for neocortical organoid polarization; however, these may vary depending on growth factor source, cell line, and protocol. From our scRNA-seq data, only a subset of candidate markers could be validated due to limited antibody availability and performance. Our organoids were analyzed up to W14, corresponding to the mid-second trimester, when areal transcriptomic signatures begin to emerge. Future studies extending beyond this time frame are needed to resolve the temporal dynamics of molecular pathways driving neocortical arealization.

## Acknowledgements

We are extremely thankful to all the lab members of the Watanabe lab for constructive feedback on our manuscript. This project has been supported by the NIH R00HD096105, NSF RECODE 2225624, New Investigator Award from the UCI School of Medicine to MW; the FRAXA Postdoctoral Fellowship to YCT; the postdoctoral training grant from the California Institute for Regenerative Medicine (CIRM; EDUC4-12822) to HO; the CIRM BRIDGES program (EDUC2-12734) to ST; the CIRM COMPASS program (EDUC5-13637) to RAG; NSF grants (DMS1763272 and CBET2134916), NIH grants (R01GM152494 and R01AR079150), and a grant from the Simons Foundation (594598) to QN; and the Simons Foundation for Autism Research grants and NIH grants (R01-MH121521 and R01-MH123922) to MJG.

## Author contributions

MW conceptualized the project, planned experiments, supervised organoid culture experiments, prepared figures, and wrote the manuscript. YCT and MJG helped write the manuscript and contributed to the conceptualization of the project. YCT, HO, KS, IF, and MJMN planned, performed, and/or analyzed experiments. YCT, HO, IF, and MJG prepared figures. ST, BKS, JKD, RAG, BDJ, GLC provided technical assistance. XW, AAA, YCT, and CN performed bioinformatic analyses for scRNA-seq under the guidance of MW, QN and MJG.

## Declaration of interests

The authors do not have any conflict of interest.

## Methods

### Animals

FVB.129P2 mice were bred and processed following protocols approved by the Institutional Animal Care and Use Committee at the University of California, Irvine. The Guidelines for the Care and Use of Laboratory Animal from the National Institutes of Health were followed.

### Human pluripotent stem cell culture

Experiments using hESC and hiPSC lines were approved by the Human Stem Cell Research Oversight Committee (hSCRO) at the University of California, Irvine. H9 hESC and Fragile X Syndrome patient iPSC lines (FX11-7, FX13-2) were obtained from the WiCell Institute, and XF iPSC line was obtained from the UCLA Broad Stem Cell Research Center Core after the material transfer agreement. The organoids used for this study were generated from hESC H9 cells between passages 60 to 75, and hiPSC FX11-7 and XF cells between passages 20-40. The stem cells were seeded with Mictomycin-C inactivated mouse embryonic feeders (PMEF-CF, Millipore Sigma) on 0.1% gelatin-coated tissue culture plate and maintained with the medium composed of DMEM/F12 (HyClone, 16777-135) with 20% knockout serum replacement (KSR, Invitrogen), non-essential amino acids (NEAAs, Invitrogen), GlutaMAX (Invitrogen), 100 mg/mL of Primocin (InvivoGen), 0.1 mM beta-mercaptoethanol (Invitrogen), and 10 ng/mL of fibroblast growth factor 2 (FGF2, Invitrogen). The medium was refreshed daily. The stem cell cultures were kept in an incubator with 5% CO_2_ at 37°C. The cells were passaged every six days by using StemPro EZ Passage tool (Invitrogen) and diluted in 1:3 - 1:7 ratios, depending on the confluency.

### Cortical organoid generation

The method for generating cortical organoids has been previously described^4, 21^. Once hESC and hiPSC colonies reached the confluency of 70-80%, they were detached with dispase (STEMCELL Technologies, 7913) and dissociated with 0.05% trypsin-EDTA (Gibco, 25300054) with Dnase 1 (Worthington, LK003172) and removed cell clumps using cell strainer of pore size 100 µm. A total of ∼9,000 cells per well were plated into low-attachment v-bottom 96-well plates (Sumitomo Bakelite, MS9096V) and aggregated by centrifuging at 290 *g* for two minutes. From day (D) 0 to 15, the organoids were cultured in the neural induction medium, consisting of Glasgow’s Minimal Essential Medium (GMEM, Gibco, 11-710-035), 20% KSR (Gibco,10828028), Non-Essential Amino Acids Solution (NEAA, Gibco, 11-140-050), 100 mg/mL of Primocin (InvivoGen, ant-pm-2), 0.1 mM b-mercaptoethanol (Gibco, 21-985-023), sodium pyruvate (Gibco, 11-360-070). We also added WNT inhibitor IWR-1-endo (IWR1e; Calbiochem, 681669) and TGF-ß inhibitor SB431542 (SB; Stemgent, 04-0010-103 µM) to the medium for cortical organoid induction, and the amount of these inhibitors depended on the cell lines: 3 µM IWR1e and 3 µM SB for H9; 1 µM IWR1e and 1 µM SB for XF; 1 µM IWR1e for FX11-7. In addition, 20 µM ROCK inhibitor Y-27632 (BioPioneer, SM-008) was supplemented in the neural differentiation medium for the first six days to reduce cell death. To evaluate the effects of the CEPT small-molecule cocktail^21^, we treated organoids with a combination of 50 nM Chroman 1 (MedChem Express, HY-15392), 5 µM Emricasan (Selleckchem, S7775), polyamine supplement (Sigma-Aldrich, P8483) diluted 1:1,000 following the manufacturer’s instructions, and 0.7 µM trans-ISRIB (Tocris, 5284). Organoids were exposed to the CEPT cocktail for either 1 or 6 days, replacing the conventional ROCK inhibitor treatment. Half of the culture medium in each well was refreshed every 2-3 days. The culture was maintained at 5% CO_2_ at 37°C. Morphological images were acquired using a BZ-X810 microscope (Keyence), and area measurements were conducted using the BZ-X800 Cell Analyzer Module. We did not observe any improvement with the CEPT cocktail compared to the ROCK inhibitor (Figure S1B).

### Cortical organoid arealization and maintenance

To anteriorize or posteriorize organoids, morphogens and small molecules were added to the medium between D15 to 21. At D15, the organoids were transferred to Petri dishes for floating culturing, and depending on the experimental group, the organoids were bathed in a medium containing either 400 ng/mL of FGF8b (Peprotech, 10025) or 50 ng/mL of BMP4 (Peprotech, 120-05ET) and 3 µM of CHIR-99021 (Tocris Bioscience, 442310). The control group received culture media with vehicles. On D18, the medium containing the morphogens and small molecules was refreshed, and the base medium was switched from the neural differentiation medium to neural maintenance medium at 37°C with 5% CO_2_ and 40% O_2_. The neural maintenance medium was comprised of DMEM/F12 (HyClone, 16777), N2 supplement (Gibco, 17502048), GlutaMAX (Gibco, 350500), chemically defined lipid concentrate (CDLC, Gibco, 11905031), primocin (InvivoGen, ant-pm-2), 5µg/mL heparin (Sigma-Aldrich, H3149), 1% matrigel (growth factor reduced, Fisher Scientific, CB-40230), and 0.4% methylcellulose (Sigma, M7140). B27 supplement without Vitamin A (Gibco, 12587010) was added to the neural maintenance medium after W5. For the Vitamin A experiment (Fig. S2C-E), the regular B27 supplement with Vitamin A (Gibco, 17504044) was used in the neural maintenance medium from W5 to W10. At W8, the organoids were transferred to Lumox dishes (Sarstedt, 94.6077.410), and human LIF (Peprotech, 300-05) was supplied in addition to the neural maintenance medium with B27 without Vitamin A. To ensure the organoids were exposed to sufficient nutrients and oxygen, the organoids were halved using microscissors at W5, W8, W10, and W12. The medium was refreshed every 2-3 days until W14.

### Single-cell RNA sequencing

We collected FGF-, no growth factor-, and BMP-treated organoids at W5, W8) and D98 (W14) from H9 (two batches for each time point) and XF (one batch for each time point) lines. For W5 samples, we pooled 25 organoids into one sample for each condition. For W8 samples, we pooled 15 organoids into one sample for each condition. For W14 samples, we pooled 15 organoids into one sample for each condition. After single-cell dissociation, a small portion of the single-cell suspensions of each sample was kept for quality control of the RNA integrity, and the rest of the single-cell suspensions were immediately processed using the Evercode™ Cell Fixation V2 kit (Parse Biosciences, ECF2001), following the manufacturer’s protocol. During the fixation procedure, the single-cell suspensions went through 40 µm cell strainers to eliminate cell clumps and were centrifuged at 200 g for 10 minutes. The samples were then cryopreserved at −80°C until all samples were ready for library preparation. The RNA quality control was conducted with Agilent 2100 Bioanalyzer, and the samples had RIN values of 9.2-10. At the UCI Genomic Research & Technology Hub, all the fixed single-cell suspensions were processed for RNA libraries with Evercode™ WT Mega V2 (Parse Biosciences, ECW02050) and sequenced all together with NovaSeq 6000 S4 flow cell, targeting 10,000 cells for a sequencing depth of 50,000 reads per cell.

### scRNA-seq data processing

Split-pipe software (Parse Biosciences, v1.0.3) was used for gene alignment, annotation and quantifying gene expression. GRCh38.p14 was used as the reference human genome assembly (Ensembl). The total number of sequenced cells was 296,056 cells in 18 samples, with the mean reads per cell around 34,131. We used Seurat “subset” function to obtain high-quality cells: <5% of total counts due to mitochondrial gene expression and 500<nFeatures<25000. After filtering, a total of 154,802 cells passed across 18 samples. Gene expression counts were normalized to have a total count of 10,000, then log-transformed, and scaled using Seurat’s “NormalizeData” with normalization.method = “LogNormalize” and “ScaleData” method. We identified the top 2000 highly variable features using Seurat “FindVariableFeatures” with selection.method = “vst”. Next, PCA is performed on the scaled data using “RunPCA”, where the top 2000 variable features are used as input. Next, cells are clustered using Seurat “FindNeighbors” based on the top 15 principal components, and then “FindClusters” with resolution 0.5. Further iterations of subclustering are then iteratively run based on known markers (see Table S1). The above functions are performed using Seurat v5.

### Pseudobulk differential gene expression analysis

To identify significant transcriptional changes due to FGF8 or BMP4/CHIR treatment, we performed differential expression analysis using edgeR (version 4.2.2)^43^. Prior to analysis, to reduce statistical noise and enable cell-type-specific analysis, for each technical replicate, we aggregated the raw, unnormalized counts across cell type using aggregateAcrossCells provided by scuttle (version 1.14.0)^44^.

We focused on analyzing differences induced by FGF8 or BMP4/CHIR treatment at specific timepoints. As we were interested in comparing not only FGF8- or BMP4/CHIR-treated conditions to the control condition, but also the FGF8 and BMP4/CHIR treatments directly, we analyzed differential expression using the design, ‘∼0 + day_condition’, where the variable day_condition is concatenated from two variables: day, which can take one of three values, ‘D35’, ‘D56’, or ‘D98’, and condition, which can be one of three values, ‘no GF’, ‘FGF8’, or ‘BMP4/CHIR’. Therefore, ‘day_condition’ can take one of nine values.

We followed standard edgeR analysis workflow to infer differentially expressed genes. First, we used filterByExpr to remove lowly expressed genes (using default parameter values). We then used calcNormFactors to estimate and then normalize the pseudobulked counts. Gene-wise dispersion values used for differential expression analysis were calculated using estimateDisp, using a negative binomial-based likelihood. We then used glmQLFit to fit a negative binomial generalized log-linear model to the normalized pseudobulked counts using quasi-likelihood methods. Finally, differentially expressed genes for each comparison—FGF8-treated vs. no GF, BMP4/CHIR-treated vs. no GF, FGF8-treated vs. BMP4/CHIR-treated— were inferred using glmQLFTest, which performs a quasi-likelihood-based F test for each gene. We defined a gene to be differentially expressed if the absolute log-fold change of its mean expression satisfied │log_2_FC│>0.5 and the false discovery rate satisfied FDR<0.05, where false discovery rates were calculated using the Benjamini-Hochberg procedure^45^.

### Cell-cell communication in scRNA-seq samples

To identify significantly active signaling pathways across FGF or BMP treatment, we used CellChat (version 2.2.0) (https://doi.org/10.1038/s41596-024-01045-4) to infer pairwise ligand-receptor interactions between cell type groups. We focused on W14 samples, analyzing communication FGF-treated and BMP-treated populations separately and only considered communication between apical radial glial cells (aRGC), intermediate progenitors (IP), deep-layer neuron-like cells (DLN), upper layer neuron-like cells (ULN), newborn neurons, and Cajal-Retzius cells (CR). Log-normalized counts and cell type annotations were used for CellChat inference.

For each condition, we followed standard CellChat analysis workflow. First, to reduce computation time, we subsetted the gene expression matrices to only consider signaling genes, using subsetData. Then, we used identifyOverExpressedGenes to determine which signal ligand and signal receptor genes were significantly expressed within cell type groups, using a permutation test. Following this, we used identifyOverExpressedInteractions to determine statistically significant ligand-receptor pairs with respect to gene expression. Ligand-receptor interaction scores were calculated using the tri-means of ligand and receptor unit gene expression in potential sender and receiver cell type groups, respectively, using computeCommunProb. To reduce bias due to signals sent or received by cell type groups with small population numbers, we removed cell types with population sizes fewer than 0.1% of the total cell number. These scores were then aggregated to the level of signaling pathways using computeCommunProbPathway. We then merged the CellChat objects for FGF and BMP-treated groups to analyze differentially incoming and outgoing signals, calculating the information flow using rankNet.

### Calculation of areal identity scores

To associate cells and cell types to their potential areal identity, for each cell, we calculated a gene score using two gene sets, one associated with frontal lobe, prefrontal cortex area identities and one associated with temporal and occipital lobes and visual areas identities (Table S6). These gene sets were manually curated from previous studies^12, 14, 15, 20^, yielding a set of 125 genes associated with anterior areal identities and a set of 48 genes associated with posterior areal identities. We calculated the cellwise gene set scores using Scanpy’s (version 1.10.1) score_genes, which calculates the relative difference between the average expression of genes within the set and a random sample of 50 genes not within the set. For all gene score calculations, we used the log-normalized gene expression values (with a pseudocount of 1).

To determine whether the gene score distributions significantly differed between conditions, we used SciPy’s (version 1.13.0)^46^ tukey_hsd to perform Tukey’s Honest Significance Distance test^47^, which tests for significant differences in distribution means between all possible pairwise combinations of groups. For each pair of conditions, we deemed the gene score distributions to be significantly different if the computed p-value satisfied p<0.05.

### Immunohistochemistry

The organoids were collected at W5, W8, and W14 and fixed in 4% paraformaldehyde/PBS solution for 20 minutes on ice, incubated in 30% sucrose/PBS solution for cryoprotection on ice, and embedded into Tissue-Tek Optimal Cutting Temperature medium (Sakura, 25608-930) in cryomolds (Sakura, 25608-924). The organoid samples were cryosectioned at 12 µm (Leica, CM3050S) and mounted onto Superfrost™ Plus Microscope Slides (Fisherbrand, 12-550-15). Immunostaining was performed using the method previously described^3, 20^. Organoid sections were incubated in the blocking buffer, consisting of 0.1% heat-inactivated horse serum (Donor equine serum, Hyclone, SH30074.02), 0.1% Triton-X, 0.01% sodium azide, and PBS, for one hour at room temperature. Primary antibodies (Table S2) were diluted in the blocking buffer and incubated overnight at 4°C. The sections were then washed three times with PBS containing 0.1% Triton X (PBST) before the incubation of secondary antibodies and Hoechst 33258 for one hour at room temperature. After washing with PBST, the sections were coverslipped with Molecular Probes™ ProLong™ Diamond Antifade Mountant (Life Technologies, P36970).

### Confocal microscopy

Images were collected (>2048 x 2048 px), using the Zeiss LSM 900 Airy Scan 2 (Zen Blue 3.3 software) mode. The same acquisition settings were used within the experimental sets for comparisons between groups. Post-acquisition adjustments, such as the brightness and contrast, were performed using ImageJ/Fiji^48^ and Adobe Photoshop, and the same adjustment settings were uniformly applied to all groups from the same experimental set. Cell number quantification was performed manually by using the Cell Counter function in ImageJ/Fiji. The percentage of cells was then calculated by normalizing to the total live cell number or specific cell types specified in the figure legends. Fluorescent intensity quantification was performed on the original CZI files without any adjustment. The region of interest was hand-circled around the periphery of the organoids or around the ventricular zone, and the raw density and integrated density were measured. The values for integrated density were used for further analysis. For CBLN2 quantification, we defined cells with much higher SATB2 expression than CTIP2, or SATB2 single-positive cells, as ULN.

### RNA isolation and quantitative real-time PCR (RT-qPCR)

The organoids were collected at W5, W8, and W14, and 4-6 organoids were pooled into one sample. The samples were lysed with the RLT buffer, and the total RNA was extracted using the RNeasy Mini Kit (Qiagen, 79216), following the manufacturer’s protocol. For cDNA synthesis, 1000-2500 ng of total RNA from each sample was used and processed with SuperScript™ IV VILO Master Mix (Invitrogen, 11756500) or SuperScript™ IV First-Strand Synthesis System (Invitrogen, 18091050). PowerTrack SYBR Green Master Mix (Applied Biosystems, A46109), primer pairs of the genes of interest (Integrated DNA Technologies), and 30ng of cDNA were used for RT-qPCR. For each gene of interest, three or more biological replicates and 2-3 technical replicates were included. The samples were loaded into 384-well plates (Invitrogen, 4309849) and processed with QuantStudio 7 Real-Time PCR System (Applied Biosystems). The relative expression level of each gene was calculated by normalizing it to GAPDH. Primers used in this study are listed in Table S3.

### Western blot

Samples from H9 were collected at W0 and W12 as positive controls. Samples from FX11-7 were collected at W0, 1.5, 2.5, 5, 8, 10, 12, and 14. In all cases, the medium was removed, and the tissues were immediately flash frozen until further processing steps. To lyse the samples, RIPA lysis buffer (50mM Tris HCl, pH8; 150 mM NaCl; 1% NP-40; 0.5% sodium deoxycholate; 0.1% SDS) with protease and phosphatase inhibitors (Thermo Scientific, 78440, 78420) was added to the samples, followed by mechanical grinding. The samples were then incubated for 10 minutes on ice, followed by pipetting with P1000 to further break down the tissue. The samples were spun down at 20,000 g for 15 minutes at 4°C. Clear supernatant was collected in a new tube. Protein quantity was determined by BCA assay (Thermo Scientific, 23227), and 30 ng of protein from each sample was used for Western blot (WB). Loading dye (BioRad, 1610747) containing β-mercaptoethanol was added to each lysate, and the mix was heated at 95°C for 5 minutes. The heated lysates and protein ladder (BioRad, 1610373) were loaded into 8–16% Mini-PROTEAN® TGX™ Precast Protein Gels (BioRad, 4561105) and the gel was run in Tris-glycine running buffer for 1 hour at 120V. The protein samples were then transferred onto a 0.45 μm nitrocellulose membrane (BioRad, 1704158) using the Trans-Blot Turbo Transfer system (BioRad, 1704150). After transfer, the membrane was incubated in the blocking solution (Nacalai, 13779-14) for 5 minutes, and incubated in the blocked membrane in primary antibody solution on a shaker at 4°C overnight. Primary antibodies used for WB were mouse anti-FMR1 (148.1) (Santa Cruz Biotechnology, sc-101048) and goat anti-GAPDH (Abcam, ab9483). After three times of TBST washes (5 minutes each), the membrane was incubated with secondary antibodies conjugated with HRP for 1.5 hours at room temperature on a shaker. The membrane was then washed three times with TBST (5 minutes each). To visualize the protein bands, we used SuperSignal West Femto Maximum Sensitivity Substrate (ThermoFisher, 34095) and Azure c600 Molecular Imager to detect the chemiluminescent signals. ImageJ was used to adjust the brightness and contrast of the entire gel.

### Statistical analysis

Specific statistical analyses are listed in the figure legends. In the cases when the FGF8 or BMP4/CHIR group was directly compared to No GFs, unpaired t-Test was used for comparing the percentage of cells, RT-qPCR data, and number of organizers (Figures 2D, 2E, and 2G). In the cases of comparing all groups together, one-way ANOVA was performed for RT-qPCR and fluorescent quantification. In the case of comparing the percentage cells in neurotypical and FXS organoids, we performed Two-way ANOVA. In the case of comparing BA4-6 and BA17 for the expression of SOX4 and SOX11, Wilcoxon signed-rank test was performed to compare the paired samples from the same individuals.

---

**Supplementary figure 1:**
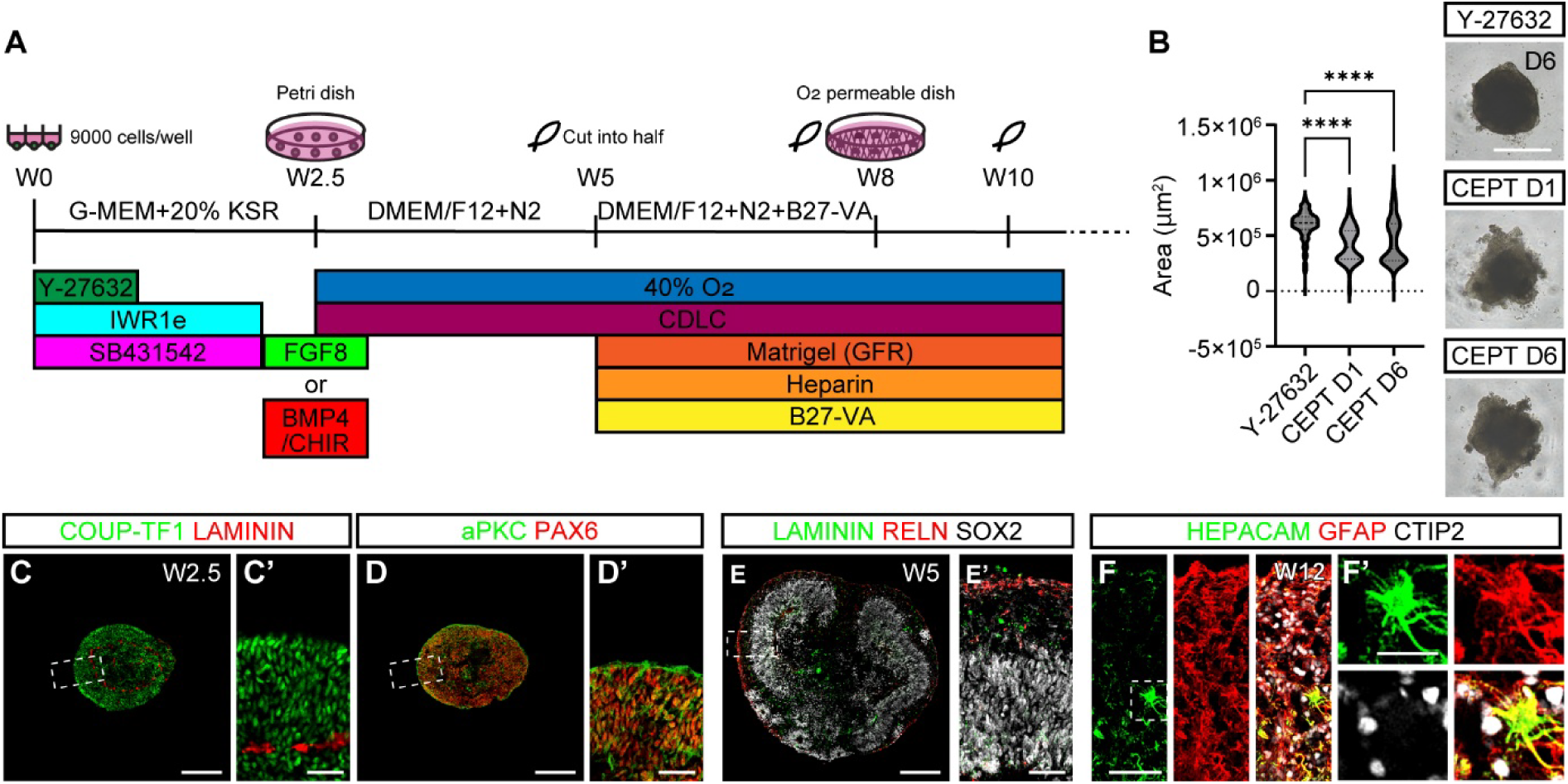
Neocortical organoid method and developmental progressions. (A) Neocortical organoid culture method. (B) Area measurement of W2.5 organoids comparing the 6-day ROCK inhibitor Y-27632, 1-day CEPT, and 6-day CEPT treatments. Scale bar: 500 µm. (C-F’) Dynamic laminar layer formation of W2.5 (C-D’) and W5 organoids (E, E’). The basement membrane is marked by LAMININ, and the apical membrane is marked by aPKC. Left panels: whole organoids. Scale bars: 200 µm. Right panels: enlarged images from the boxed region. Scale bars: 50 µm. (F) Astrocyte formation in W12 neocortical organoids, labeled with HEPACAM and GFAP. Scale bar: 50 µm. (F’) Zoom-in images from (F), white dotted square. Scale bar: 20 µm.

**Supplementary figure 2:**
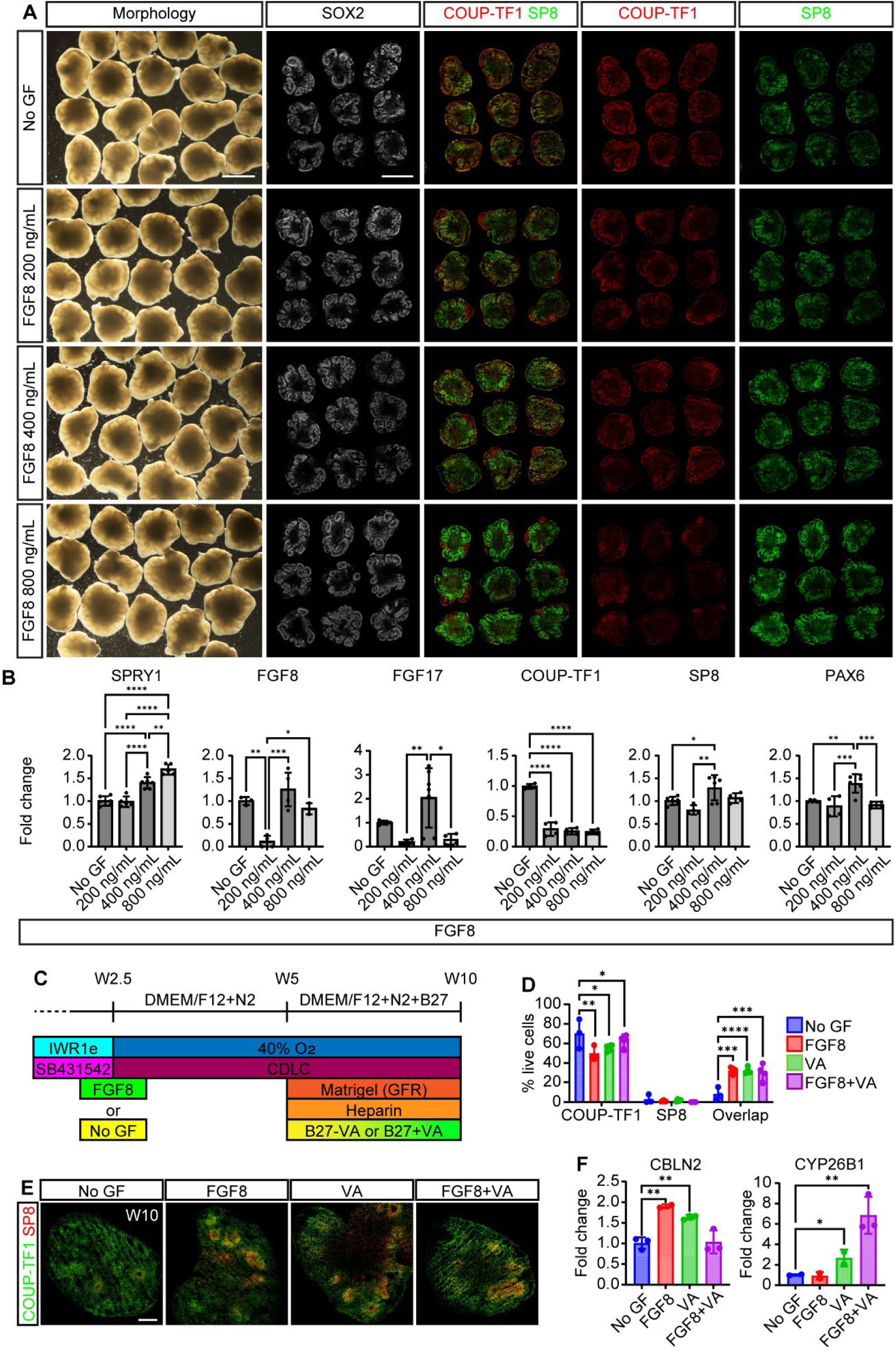
FGF8 titrations and Vitamin A application to anteriorize neocortical organoids. (A) Brightfield (left panel) and immunostaining of anterior- and posterior-high transcription factors, SP8 and COUP-TF1, respectively, in neocortical organoids treated with no GF, 200 ng/mL, 400 ng/mL, and 800 ng/mL of FGF8. Scale bars: 1000 µm. (B) Gene expression levels in organoids treated with varying FGF concentrations measured by RT-qPCR. FGF direct target gene, SPRY1. FGF growth factors, FGF8 and FGF17. Canonical anterior transcription factors, SP8 and PAX6. A canonical posterior transcription factor, COUP-TF1. N=4-6 from 2-3 batches/condition. Statistics: One-way ANOVA with post hoc test Tukey. Data are presented as mean ± SD. *p<0.05; **p<0.01; ***p<0.001; ****p<0.0001. (C-F) Comparison of the anteriorization effect in four conditions: no GF-treated, FGF8-treated, Vitamin A-treated, and FGF8-/Vitamin A-treated organoids. (C) Experimental paradigm. (D) Percentage of COUP-TF1^+^, SP8^+^, and double-positive populations in four conditions. N=3-4/condition. Statistics: Two-way ANOVA with post hoc test Tukey. Data are presented as mean ± SD. *p<0.05; **p<0.01; ***p<0.001; ****p<0.0001. (E) Representative immunochemical images of W10-organoids from 4 conditions, stained with COUP-TF1 and SP8. Scale bar: 200 µm. (F) RT-qPCR results of CBLN2 and CYP26B1 expressions from four conditions. N=2-3/condition. Statistics: Two-way ANOVA with post hoc test Tukey. Data are presented as mean ± SD. *p<0.05; **p<0.01.

**Supplementary figure 3:**
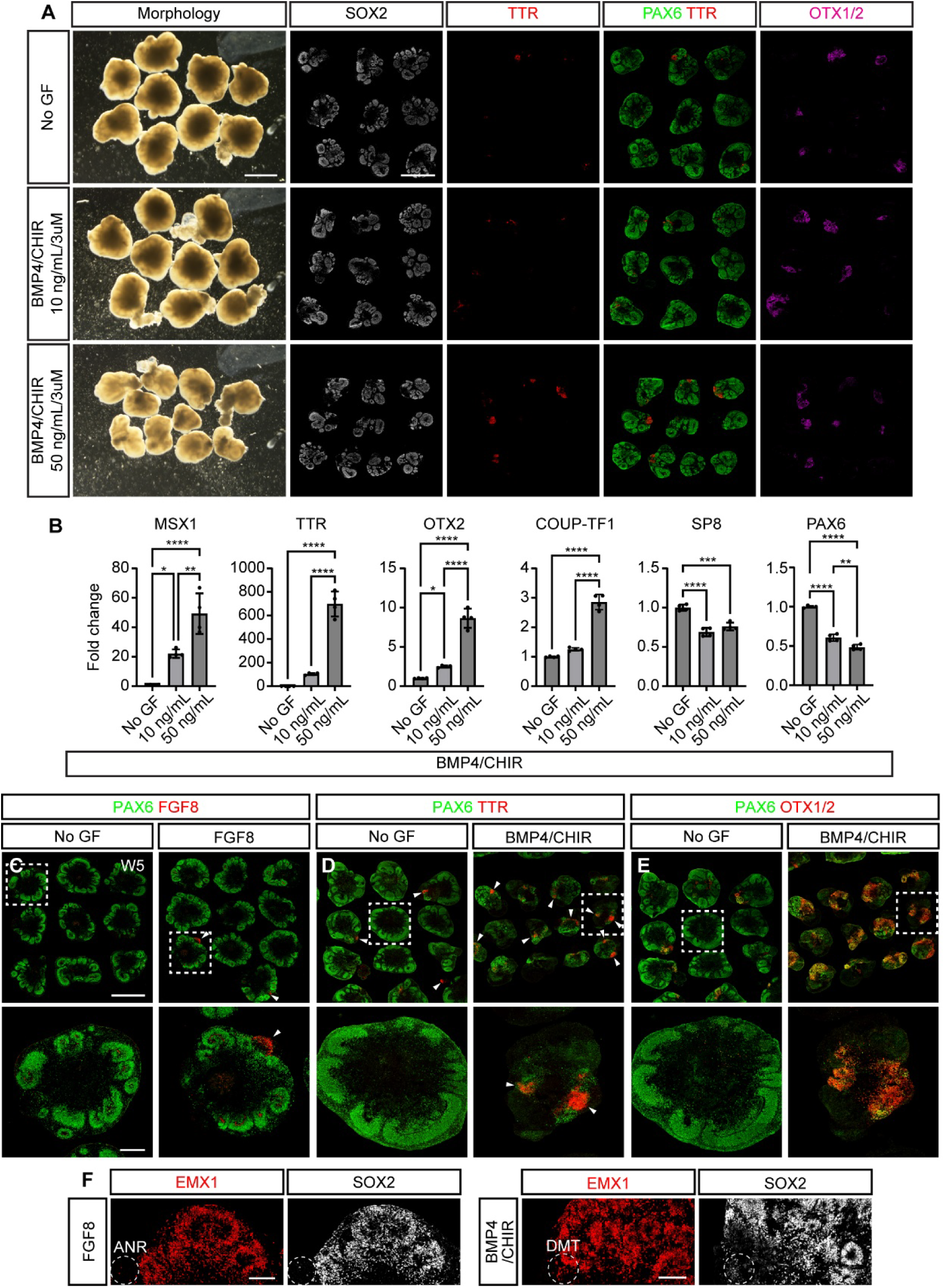
BMP4 titrations with 3µM of CHIR-99021 to posteriorize neocortical organoids and the organizer formation with FGF8 or BMP4/CHIR application. (A) Brightfield (the most left panels) and immunostaining images of DMT markers (TTR and OTX1/2) and generic neural progenitor markers (SOX2 and PAX6) in neocortical organoids treated with no GF, 10 ng/mL, and 50 ng/mL of BMP4. Scale bars: 1000 µm. (B) Gene expression levels in organoids treated with various BMP concentrations, measured by RT-qPCR BMP direct target gene, MSX1. DMT markers, TTR and OTX2. Canonical anterior-high transcription factors, SP8 and PAX6. Canonical posterior-high marker, COUP-TF1. N=4 from two batches/condition. Statistics: One-way ANOVA with post hoc test Tukey. Data are presented as mean ± SD. *p<0.05; **p<0.01; ***p<0.001; ****p<0.0001. (C) The anterior neural ridge (ANR) formation is marked by FGF8 expression in organoids treated with FGF8 (400 ng/ml). Upper panels: Tile images include multiple organoids from one batch. Scale bar: 1000 µm. Lower panels: Example organoids from the white squares in the tile images. Scale bar: 200 µm. (D-E) The posterior organizer, DMT regions marked by TTR^+^ (D) and OTX1/2+ (E) in organoids treated with no GFs and BMP4 (50ng/ml) and CHIR-99021 (3µM). Upper panels: Tile images include multiple organoids from one batch. Scale bar: 1000µm. Lower panels: Example organoids from the white squares in the tile images. Scale bar: 200µm. (F) Cortical marker EMX1 immunostaining of FGF8-treated and BMP4-treated organoids to ensure all analyses were done in the EMX1^+^ neocortical region. Scale bar: 100µm.

**Supplementary figure 4:**
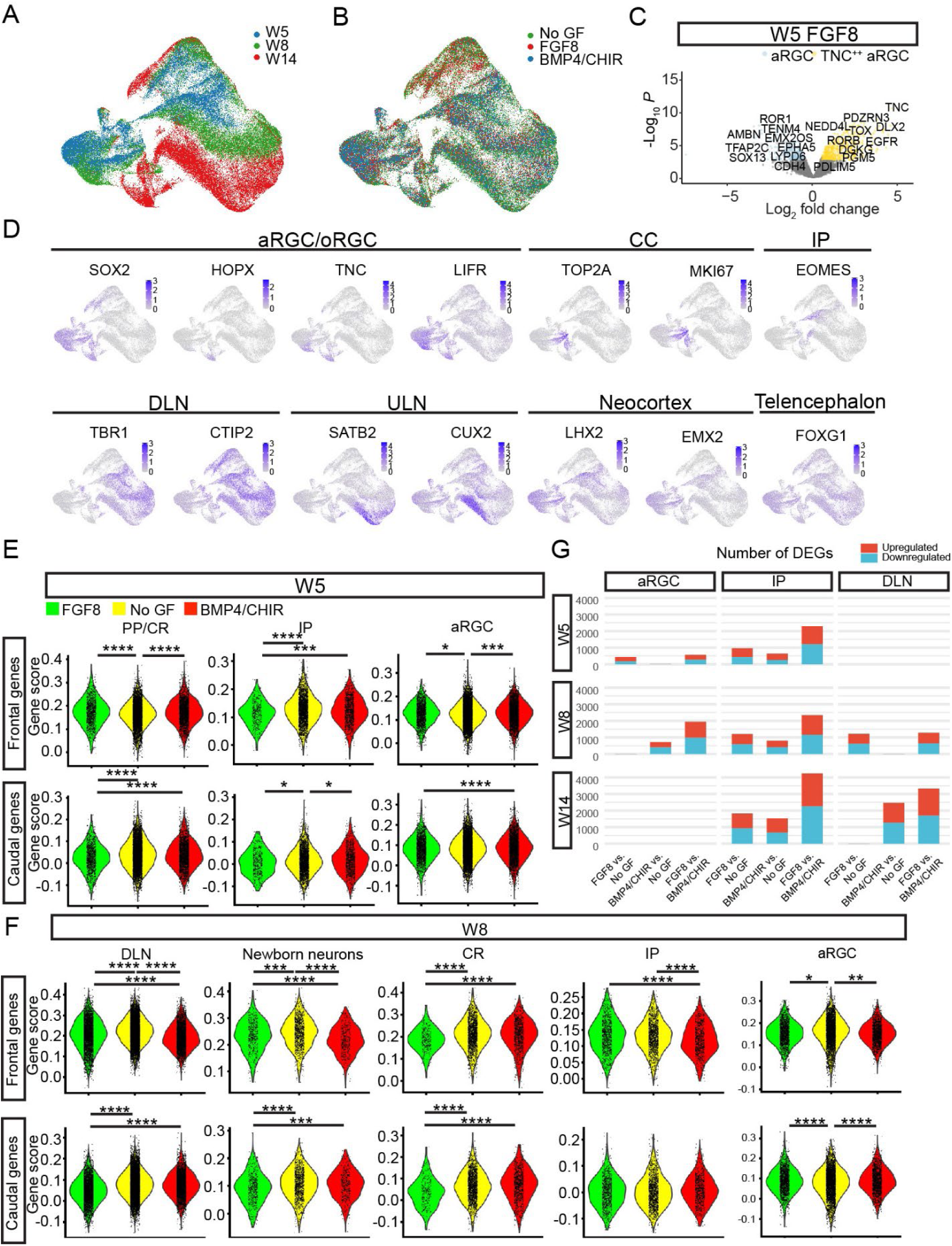
Extended data for Figure 3. Neocortical organoids reveal temporally dynamic and cell-type-specific gene signatures associated with frontal and caudal areas. (A-B) UMAP of the scRNA-seq data color-coded by age (A) and by conditions (B). (C) Volcano plot comparing aRGC and TNC^++^ aRGC in W5-organoids treated with FGF8 to show DLX2 and TOX upregulation and EMX2 downregulation in TNC^++^ aRGC, suggesting its ganglionic eminence aRGC identity. Colored: genes with FDR<0.05 and │log2FC│>0.5. Example genes are labeled. (D) Gene expression levels for canonical markers of different cell clusters are plotted on the UMAP. (E-F) Frontal (upper panels) and caudal area (lower panels) gene scores in an individual cell at W5 (E) and W8 (F). Each cell type is plotted, comparing organoids treated with FGF8, No GFs, and BMP4/CHIR. Statistics: Tukey’s Honest Significance Distance test. *p<0.05; **p<0.01; ***p<0.001; ****p<0.0001. (G) Number of DGEs from pseudobulk analysis, comparing FGF8 vs No GF, BMP4/CHIR vs No GF, and FGF8 vs BMP4/CHIR.

**Supplementary figure 5:**
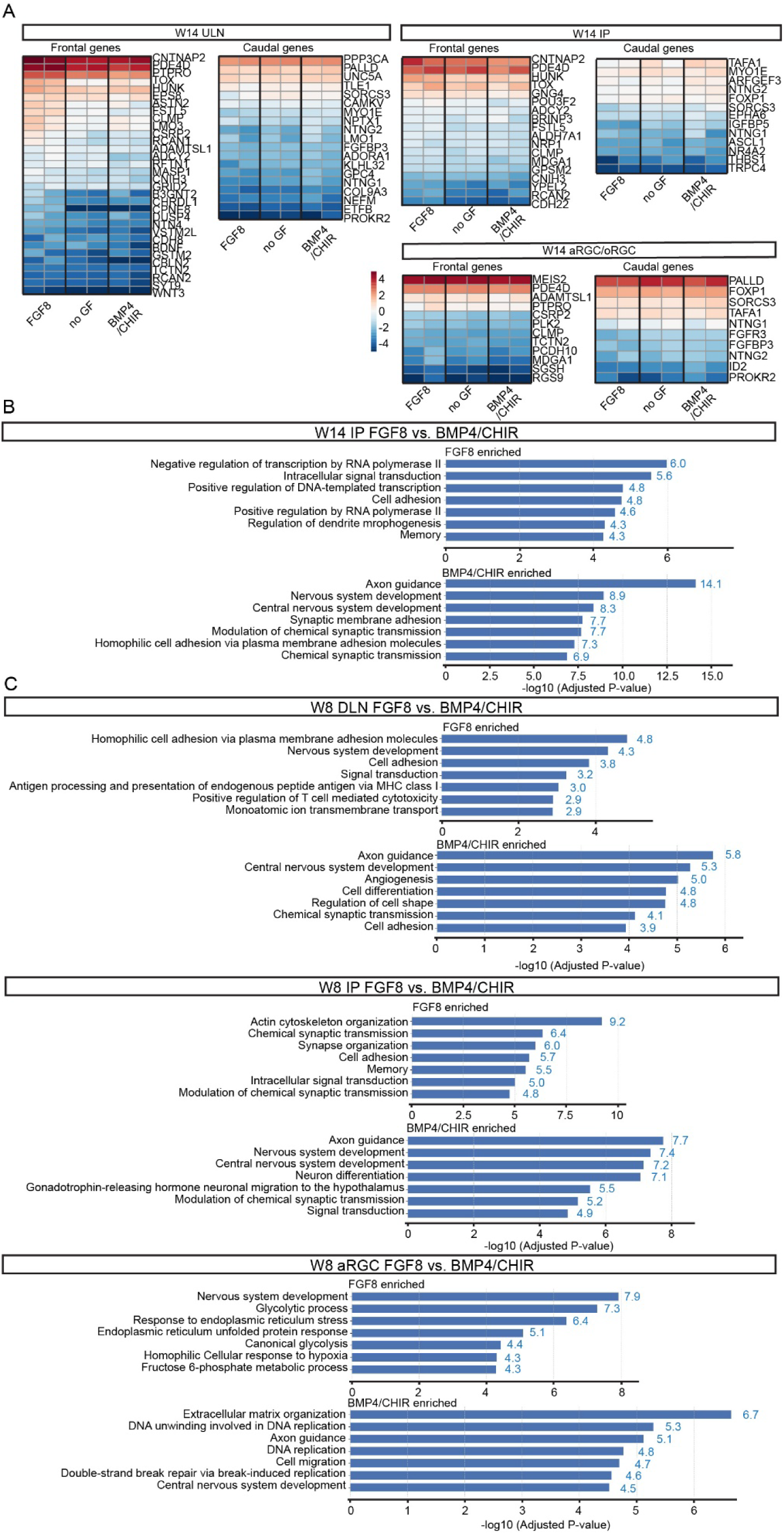
Extended data for Figure 3: gradient expression of frontal and caudal area genes and gene ontology analysis. (A) Log-normalized gene expression of known frontal and caudal area genes in W14 ULN, IP, and aRGC/oRGC. Two biological replicates for each condition: FGF8, no GF, and BMP4/CHIR. (B-C) Gene ontology analysis, using the differentially expressed genes comparing FGF8- to BMP4/CHIR-treated organoids for each cell cluster: IP at W14 (B), DLN, IP, and aRGC at W8 (C). The top seven biological processes were selected.

**Supplementary figure 6:**
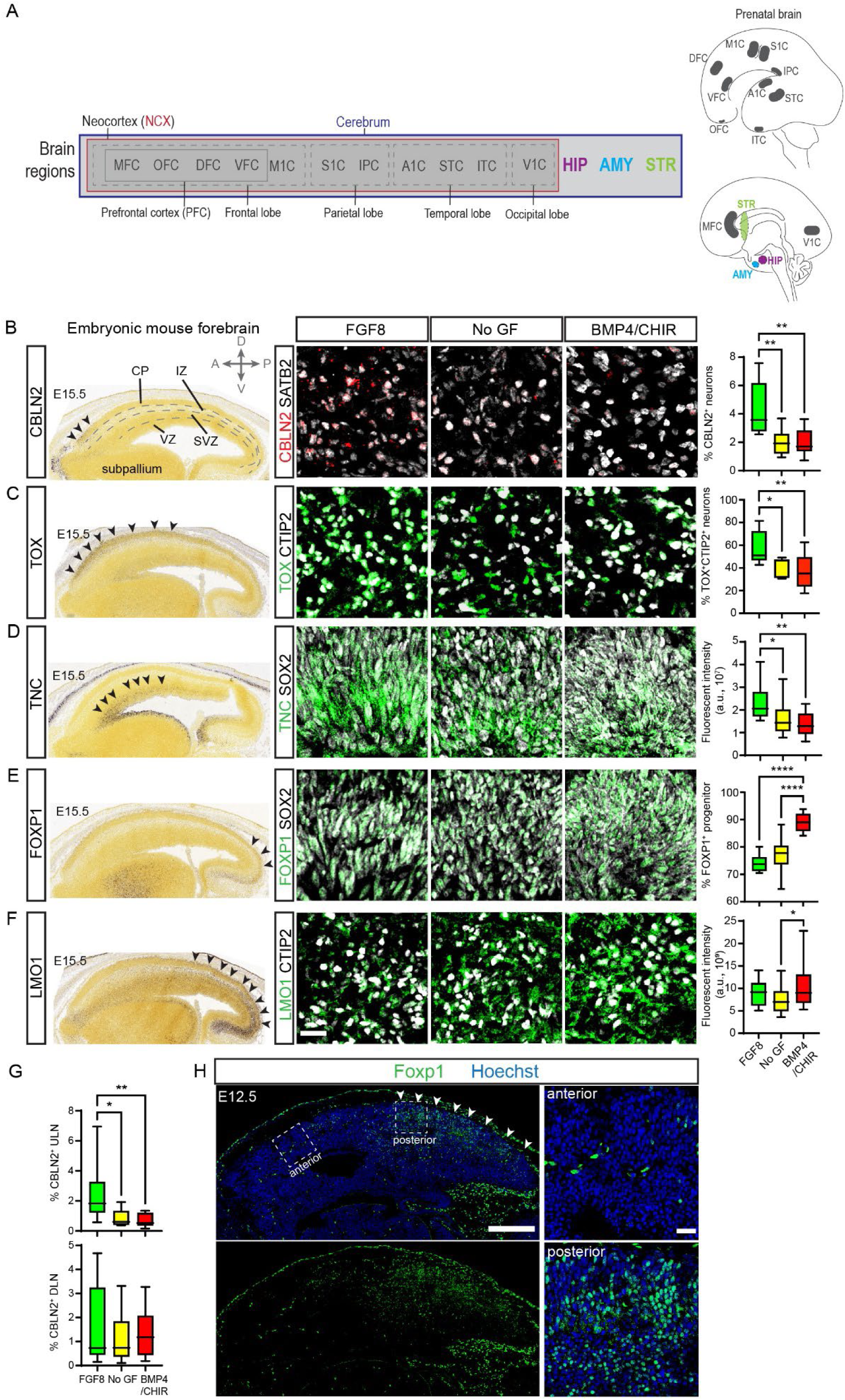
CBLN2 positive populations in ULN and DLN. (A) Schematic illustration of cortical and brain regions in the embryonic developmental brain. (B-F) Left panels: Expression patterns of Cbln2, Tox, Tnc, Foxp1, and Lmo1 by in situ hybridization of embryonic mouse cortices, from Allen Brain Atlas (developingmouse.brain-map.org). Arrowheads highlight the gene expression. Cortical layers are labeled in panel B. CP: cortical plate; IZ: intermediate zone; SVZ: subventricular zone; VZ: ventricular zone. (B) CBLN2 expression in W14 neocortical organoids. Quantification (right panel): percentage of CBLN2^+^ neurons in total (SATB2+ and CTIP2+) neurons. N = 9 organoids per condition from a total of 3 batches. (C) TOX expression in W14 neocortical organoids. Quantification (right panel): percentage of TOX^+^CTIP2^+^ neurons in total CTIP2^+^ neurons. 8-9 organoids per condition from a total of 3 batches. (D) TNC expression in W8 neocortical organoids. Quantification (right panel): fluorescent intensity in the SOX2^+^ VZ region. 12 organoids per condition from a total of 3 batches. “IntDen” values, the product of area and mean gray value, were used for comparisons. (E) FOXP1 expression in W8 neocortical organoids. Quantification (right panel): percentage of FOXP1^+^SOX2^+^ progenitors in total SOX2^+^ progenitors. 9 organoids per condition from a total of 3 batches. (F) LMO1 expression in W14 neocortical organoids. Quantification (right panel): Fluorescent intensity of whole organoids. 10 organoids per condition from a total of 3 batches. “IntDen” values, the product of area and mean gray value, were used for comparisons. Scale bar: 20 µm. Statistics: One-way ANOVA with post hoc test Tukey. *p<0.05; **p<0.01; ***p<0.001; ****p<0.0001. Data are represented as mean ± SD. (G) Percentage of CBLN2^+^SATB2^+^ ULN (left) and CBLN2^+^CTIP2^+^ DLN in total (SATB2^+^ and CTIP2^+^) neurons. N = 9 organoids per condition from a total of 3 batches. Statistics: One-way ANOVA with post hoc test Tukey. Data are presented as mean ± SD. *p<0.05; **p<0.01; ***p<0.001; ****p<0.0001. (H) Sagittal section of E12.5 mouse brain immunostained with Foxp1. Arrowheads highlight Foxp1 expression. Scale bar: 250 µm, and 25 µm for zoom-in images on the right panels.

**Supplementary figure 7:**
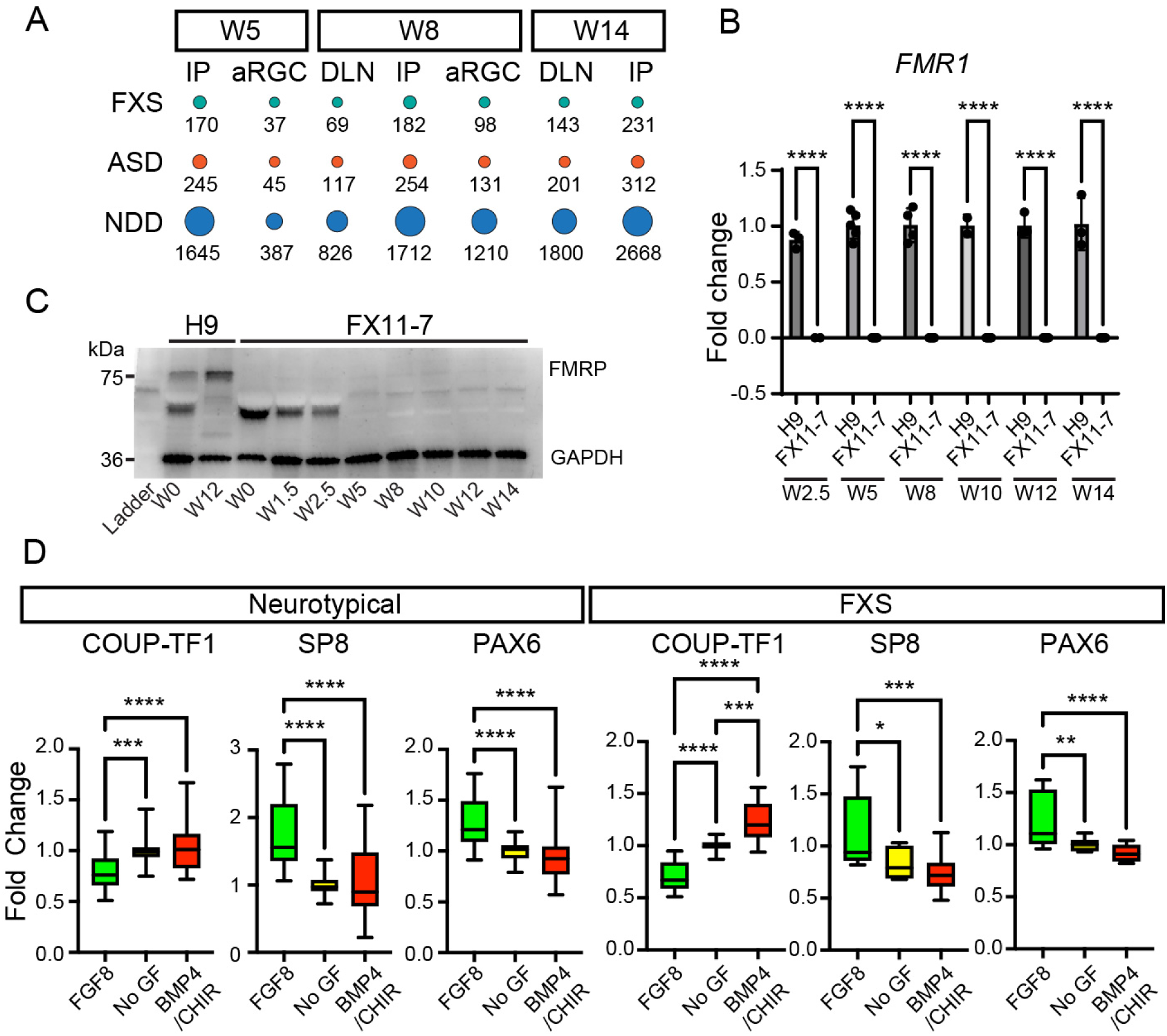
Extended data for FXS modeling. (A) Association analysis showing the number of DEGs (FGF8 vs BMP4/CHIR) from the respective cell types and time points overlapped with Fragile X Syndrome (Darnell et al, 2011)^32^, autism spectrum disorders (SFARI, https://gene.sfari.org/)^33^, and neurodevelopmental disorders (Fu et al, 2022)^34^. (B) FMR1 gene expression in H9 (control) and FX11-7 (FXS iPSC-derived) organoids from W2.5 to W14 by RT-qPCR assays. (C) Western blot showing FMRP (∼75 kDa) expression in FX11-7 organoids from W0 to W14. H9 organoids at W0 and W12 were used as positive controls. (D) Gene expression levels of COUP-TF1, SP8, and PAX6 in neurotypical control and FXS neocortical organoids treated with FGF8, no GF, or BMP4/CHIR. Neurotypical: N=30-33, including technical and biological replicates, from H9 and XF, two batches each. FXS: N=7-17, including technical and biological replicates, from FX11-7, four batches. Statistics: One-way ANOVA with post hoc test Tukey. Data are presented as mean ± SD. *p<0.05; **p<0.01; ***p<0.001; ****p<0.0001.

## Notes

### Competing Interest Statement

The authors have declared no competing interest.

